# Linking Fungal and Bacterial Proliferation to Microbiologically Influenced Corrosion in B20 Biodiesel Storage Tanks

**DOI:** 10.1101/399428

**Authors:** Blake W. Stamps, Caitlin L. Bojanowski, Carrie A. Drake, Heather S. Nunn, Pamela F. Lloyd, James G. Floyd, Katelyn A. Berberich, Abby R. Neal, Wendy J. Crookes-Goodson, Bradley S. Stevenson

## Abstract

Biodiesel is a renewable substitute, or extender, for petroleum diesel that is composed of a mixture of fatty acid methyl esters (FAME) derived from plant and animal fats. Ultra-low sulfur diesel (ULSD) blended with up to 20% FAME can be used interchangeably with ULSD, is compatible with existing infrastructure, but is also more susceptible to biodegradation. Microbial proliferation and fuel degradation in biodiesel blends has not been directly linked *in situ* to microbiologically influenced corrosion. We, therefore, conducted a yearlong study of B20 storage tanks in operation at two locations, identified the microorganisms responsible for observed fuel fouling and degradation, and measured *in situ* corrosion. The bacterial populations were more diverse than the fungal populations, and largely unique to each location. The bacterial populations included members of the *Acetobacteraceae, Clostridiaceae*, and Proteobacteria. The abundant Eukaryotes at both locations consisted of the same taxa, including a filamentous fungus within the family *Trichocomaceae*, and the *Saccharomycetaceae* family of yeasts. Increases in the absolute and relative abundances of the *Trichocomaceae* were correlated with significant, visible fouling and pitting corrosion. This study identified the relationship between recurrent fouling of B20 with increased rates of corrosion, largely at the bottom of the sampled storage tanks.

## Introduction

The use of renewable fuels is on the rise worldwide (U.S. Energy Administration (2015)). One renewable fuel, biodiesel, is composed of fatty acid methyl esters (FAME) derived from plant and animal lipids used as a replacement or additive to petroleum diesel^1^. Up to 5% v/v biodiesel is commonly used as an additive to ultra-low sulfur diesel (ULSD) to restore lubricity lost due to the removal of sulfur compounds^2,3^. Blends containing up to 20% v/v biodiesel (i.e., B20 biodiesel) can be used without changes to existing engines or infrastructure^4^. The consumption of biodiesel (including B20 biodiesel) has significantly increased worldwide over the past 10 years due to similar combustion properties to ULSD^5,6^, performance advantages, and the adoption of stricter regulation on emissions^7^.

Despite the advantages offered by blending biodiesel with ULSD, biodiesel can negatively affect fuel stability^8^. Biodiesel contains more dissolved oxygen than ULSD, reducing oxidative stability and increasing its biodegradability^9^. Biodiesel is also more hygroscopic than ULSD^10^, causing B20 to absorb and retain more water. Water is essential for microbial metabolism and growth; water entrained within fuel enables microbiological colonization of the fuel and fouling^11-14^. As microorganisms grow and foul a storage tank, conditions may become more conducive for corrosion within the storage tank^15^.

The metabolism of FAME by microorganisms in a B20 storage tank generates organic acids^15,16^ that may result in microbiologically influenced corrosion (MIC) of steel tanks or tank components. Also, exposure of polymer tank seals and fittings to organic acids can decrease their ductility and strength^17^. In addition to organic acids, production of CO_2_ during the microbial metabolism of FAME could also increase the rate of corrosion within a storage tank^18^. General production and dissolution of acid and CO_2_ within a fluid can result in uniform corrosion, where the loss of material is distributed across a surface. The accumulation of biofilms, however, can result in the localized production and concentration of corrosive compounds, the formation of galvanic couples, or the ennoblement of steel surfaces. The result is localized corrosion that is characterized by deep, penetrating pits on metallic surfaces that represent a great risk to storage tank infrastructure ^19,20^.

For over a decade, the United States Air Force (USAF) has been storing B20 biodiesel and using it in non-tactical support vehicles as a component of their Strategic Energy Plan^7,21^. The adoption of B20 biodiesel by the USAF, however, was correlated with an increase in reports of “bad fuel”, or fouling; fuel was reported to contain particulates that clogged fuel dispensing filters. Both fuel fouling and MIC have the potential to impact the operation of B20 biodiesel storage and distribution systems. The link between biodegradation and corrosion has been studied in other petroleum environments such as production, drilling^22,23^, or storage of petroleum products including biodiesel^15,24^. Pure cultures or enrichments of microorganisms have also been shown to degrade biodiesel blends (including B20) and induce corrosion under laboratory conditions ^13,25,26^ but studies that address the microbial community and corrosion risk within active storage tanks are currently nonexistent. To this end, we were able to conduct a rare longitudinal study, *in situ,* of microbial community dynamics linked to the corrosion of steel in B20 biodiesel underground storage tanks (USTs). Two USAF locations housing USTs with recurrent fuel quality issues were selected for this study. We hypothesized that the reported issues were the result of microbial contamination and predicted that corrosion would be greatest in USTs with the most biomass. We subsequently conducted a one-year survey of the microbial communities present at both storage locations to link fouling, fuel degradation, and MIC *in situ* within each UST.

## Methods

### Sampling and In Situ Analyses of Witness Coupons

The two USAF bases chosen for our yearlong study were located in the southeast (SE) and southwest (SW) United States. The study focused on three USTs containing B20 biodiesel at each base for a total of 6 surveyed USTs. All tanks were operated normally throughout the study, except for SE 3, which was removed from the study after 9 months due to severe microbiological contamination that required mitigation. The three tanks at SW were constructed of uncoated carbon steel and installed in the early 1950s. At the SE site, two of the tanks (SE 3, SE 4) were fiberglass and located at the same fueling station. The third tank (SE E) was made of carbon steel, lined on the outside with fiberglass, and located at another fueling station. Each of the six tanks had a large maintenance hatch (i.e. “manway”) that was used as an access point for fuel sampling (SI Figure 1). This hatch also served as an attachment point for suspension of a poly-vinyl chloride (PVC) rack that held four types of materials typical of fuel systems: steel, epoxy-coated steel, and fluorocarbon and Viton™ polymers. One PVC rack was placed in each tank through the maintenance hatch and allowed to rest at the bottom of the tank. Witness coupons were removed at ≈3-month intervals at SE, and ≈6-month intervals at SW. Fuel samples were taken at each time point from the bottom of the tank via the access hatch using a 500 mL Bacon bomb fuel sampler (Thermo Fisher Scientific, Hampton, NH). As a control, sterile, uncoated witness coupons and sterile O-rings were incubated separately in 0.22 µm filter-sterilized B20 taken from fuel samples received at each site prior to exposure to USTs (n = 9 for each location). Additional information related to the construction of the sampling rack, and epoxy coating composition can be found in supplemental methods.

**Figure 1.**
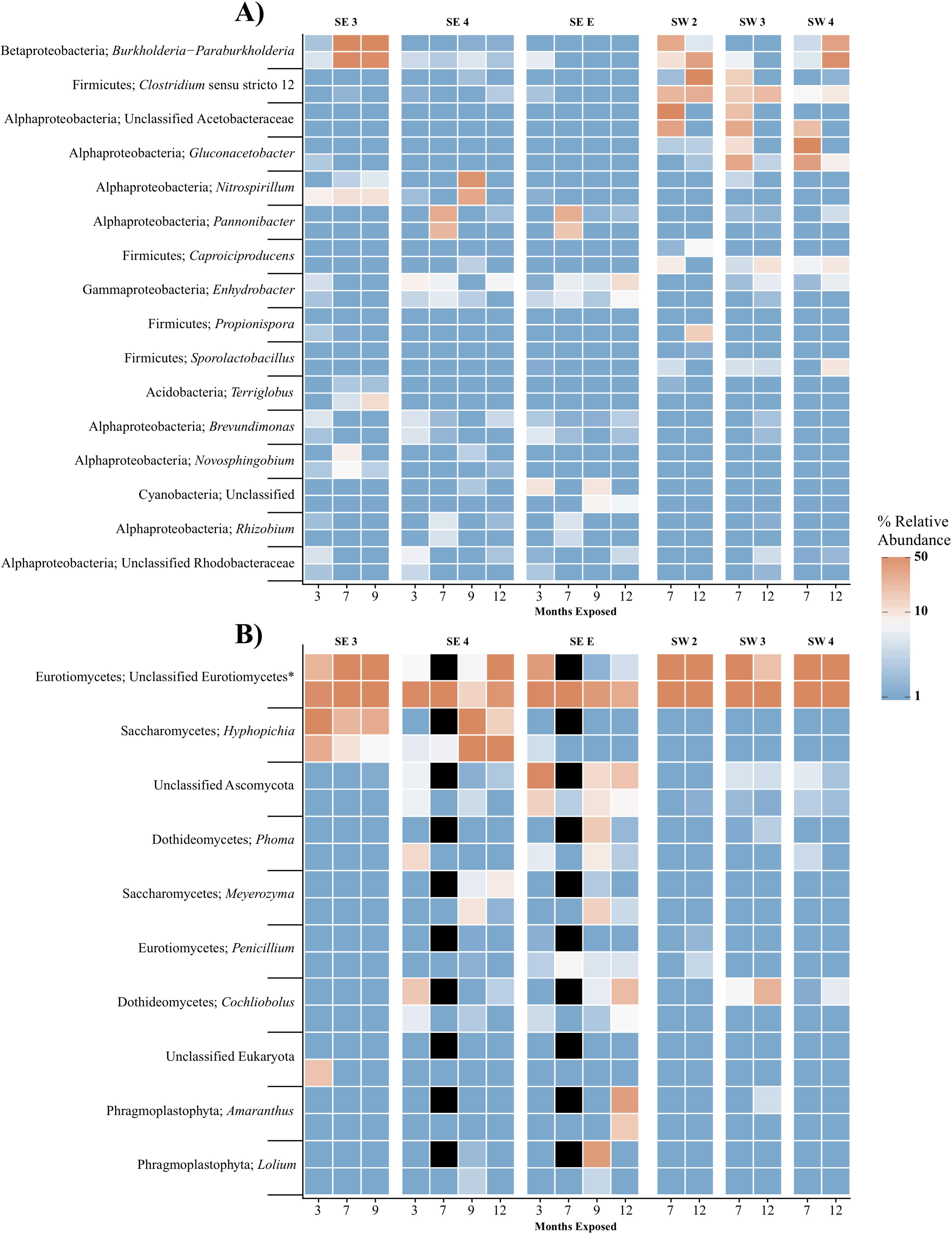
Heat map showing the relative abundance of the 20 most abundant bacterial (A) and the 10 most abundant Eukaryotic (B) OTUs within the microbial communities found in biofilms within each tank over time. For each taxon shown, the heat map is divided into samples which were suspended from the top and bottom of the PVC rack. Taxonomic identity for each OTU is shown as the most likely genus (if identifiable) and phylum. For clarity, Proteobacterial classes are shown in place of the phylum designation. Black boxes denote that insufficient numbers of sequence were avalible to analyze for those timepoints.

Replicate witness coupons (n ≥5) were removed from tanks at specified time intervals and photographed on-site. These coupons were subjected to several types of analyses: characterization of biofilm microbial communities by DNA sequencing, microscopy, and quantitation of uniform and pitting corrosion by mass loss and microscopy, respectively. A schematic representation of the sampling workflow is shown in supplemental figure S2, and further detail on the provenance of sample specimens, including polymeric materials, is available in supplemental methods. Biofilms were sampled for molecular community analyses from triplicate witness coupons immediately following their removal from the tank. Otherwise, coupons were stored in a desiccating environment prior to further analyses.

### Small Subunit rRNA Gene Library Preparation and Sequencing

Immediately upon removal of coupons from the tanks, the surface was sampled using nylon flocked swabs (Therapak Corp, Los Angeles, CA), placed into a 2.0 mL ZR BashingBead™ Lysis Tube (Zymo Research Corp., Irvine, CA) containing 0.7 mL (dry volume) of 0.5 mm ZR BashingBead™ lysis matrix (Zymo Research Corp.) and 750 µL Xpedition™ Lysis/Stabilization Solution (Zymo Research Corp.) and homogenized for 30 sec on site using a sample cup attached to a cordless reciprocating saw. Fuel samples were taken and filtered on-site, with approximately 1 L of fuel filtered through a 120 mm 0.22 µm polyether sulfone bottle-top filter. At SE, the filtered fuel was also retained for acid-index determination. The filter was quartered after sampling using a sterile scalpel. Three of these quarters were placed into individual ZR BashingBead Lysis tubes, for a total of 3 technical replicates per sampling. Samples were transported overnight at room temperature and stored at −20 °C upon receipt. Prior to DNA extraction, the samples were homogenized for an additional 30 s using a BioSpec Mini-BeadBeater-8 (Biospec Products Inc., Bartlesville, OK). DNA extractions were performed per manufacturer specifications using the Zymo Xpedition™ Kit (Zymo Research Corp.).

Libraries of bacterial, archaeal, and eukaryotic small subunit (SSU) ribosomal RNA (rRNA) gene fragments were amplified from each DNA extraction using PCR with primers that spanned the SSU rRNA gene V4/V5 hypervariable regions between position 515 and 926 (*E. coli* numbering) as described previously^27,28^. To mitigate the effects of contaminating DNA^29^, multiple extraction blanks and negative controls were sequenced from each batch of extractions. Subsequently, SSU rRNA gene libraries were sequenced using Illumina MiSeq V2 PE250 chemistry.

### Analysis of SSU rRNA Gene Sequencing Libraries

Initial quality control, demultiplexing, and operational taxonomic unit (OTU) clustering at 97 percent sequence similarity were performed as previously described^30^, with the modification of using mothur^31^ to assign taxonomy against the SILVA database (r128)^32^ formatted for use with mothur. Statistical analyses and figures were generated within R using Phyloseq^33^ and AmpVis^34^. Differences in community composition were estimated using the weighted UniFrac index^35^. The effect of site, tank, material, and the location of materials within each tank (top or bottom) were tested by a PERMANOVA within the R package Vegan^36^. Sequencing reads are available under the accessions SRR5826605-SRR5826609.

### Determination of Fuel Acid Index

The acid index of fuel samples taken from the bottom of tanks at the SE location was measured by acid titration using the ASTM standard D974^37^ method at the 3, 7, and 9 month time points. Approximately 20 g of B20 suspended in 100 mL of titration solvent (100:1: 99 Toluene/Water/Isopropyl alcohol) and 0.5mL of an indicator solution was titrated using a 0.1 N solution of KOH dissolved in isopropyl alcohol (Sigma Aldrich).

### Estimation of final coupon biomass

The growth of biofilms within the USTs was determined by measuring the biomass attached to the polymer-coated witness coupons. After 12 months of exposure, all pre-weighed polymer coated coupons were removed from all tanks at both locations, photographed on site, placed into sterile 50 mL conical tubes, and shipped overnight for further analysis. Upon arrival, all coupons were desiccated to remove any water or fuel, weighed, and then cleaned using the ASTM standard G1-03 hydrochloric acid method^38^. Biomass at 12 months was estimated as the difference in the weight of the coupons prior to incubation *in situ* and after cleaning. No degradation of the polymer coating was observed before or after removal of the biofilm from the coupon.

### Microscopy and Determination of Corrosion on Witness Coupons

For determination of general corrosion, biofilm and corrosion products were removed from coupons via ASTM standard G1-03 C.3.5^38^. After cleaning, coupons were weighed on an analytical balance, masses recorded, and compared against pre-exposure masses of each coupon. Mass loss was calculated as the difference between weight prior to incubation (initial weight) and the weight of the coupon after cleaning. A subset of steel witness coupons (n=2 per time point) was imaged by several complementary approaches: scanning electron microscopy (SEM), confocal microscopies, and white light profilometry. Coupons with intact biofilms that were intended for microscopic analysis were scored with a 2 × 14 grid to enable correlation between locations imaged (each quadrant roughly 9 × 11mm, Figure S2). Additional information on the scribing of coupons is available in supplemental methods. Select coupons were then imaged with the biofilm in place using a VHX 2000E (Keyence Corp., Itasca, IL) digital microscope from 20- 200X magnification. Areas of interest were subsequently imaged using an FEI Quanta 600 environmental scanning electron microscope (ESEM, Thermo Fisher Scientific Co., Waltham, MA). Three to five areas were imaged using the SEM per coupon.

Coupons designated for pitting measurement after cleaning were imaged using a VK-X250 3D Laser Scanning Confocal Microscope (Keyence Corp., Itasca, IL) to measure the depth of each pit depth and surface roughness. Each coupon was imaged under 10X magnification at 42 locations chosen randomly for the first coupon analyzed, and then used for all other coupons. Maximum pitting rates (Recorded as mils per year or MPY) were calculated for each sample point using the formula 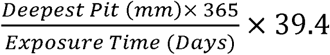 where 39.4 is the conversion factor to go from mm/year to MPY. using A pit was defined as a region that was a minimum of 10 µm below the mathematically determined reference plane (surface) and at least 200 µm^2^ in area. A Shapiro-Wilk^39^ test of normalcy (p > 0.05) was used to determine normality of the dataset, after which a non-parametric Van der Waerden test^40^ was used to compare field coupons to sterile controls. Multiple pairwise tests comparing *in situ* incubations to controls were carried out *post hoc* using a Conover test^41^ within the package PMCMR, and *p* values were corrected using the false discovery rate (FDR) method^42^ within PMCMR. To determine if there was a correlation between the presence of a biofilm and corrosion under the biofilm, areas imaged by SEM were also imaged by the VK-X250 confocal microscope and compared. Additional information on imaging methods can be found in supplemental methods.

### Results

## Microbial diversity and taxonomic composition results

Swabs taken at each time point at each UST from coated coupons, uncoated coupons, and O-rings (representing biofilm communities), and filter quarters (representing fuel communities) were sequenced after sampling. After clustering and quality control, 3.04 million reads remained, which clustered into 759 OTUs across all three domains of life from 306 samples (Supplemental table 1). Mean library size for Bacteria and Archaea was 6857 reads, and 3470 for the Eukarya (Supplemental table 1).

**Table 1.** Acid index of select fuels from SE at the 3, 7, and 9 month time points^a^

|  | 3 Months | 7 Months | 9 Months |
| --- | --- | --- | --- |
| **SE 3** | 0.17 | 0.30 | **1.51** |
| **SE 4** | 0.08 | 0.16 | 0.28 |
| **SE E** | 0.22 | 0.09 | 0.24 |
| **Receipt<sup>a</sup>** |  |  | 0.06 |

<sup>a</sup> Values are expressed in mg KOH/g B20. Bold values indicate fuel exceeds ASTM standard for acidity.
<sup>b</sup>Receipt was fuel taken directly from a delivery truck before exposure to a storage tank.

The community structure and composition of sampled biofilms varied significantly between bases (Bacteria p = 0.001 R^2^ = 0.12, Eukaryota p = 0.001 R^2^= 0.23), and between different tanks at each base (Bacteria p = 0.001 R^2^ = 0.37, Eukaryota p = 0.001 R^2^ = 0.34). Members of the Acetobacteraceae (acetic acid bacteria) or the Clostridiaceae group 1 were the most abundant taxa detected in the tanks at SW, whereas members of the Rhodospirillaceae and Sphingomonadaceae were more abundant in SE tanks (Figure 1a). An OTU (OTU 1 within the Eukaryotic dataset) that was present and often very abundant at both locations was identified as an unclassified member of the Eurotiomycetes. Additional identification via BLAST suggested that it was a filamentous fungus, most likely a member of the genus *Byssochlamys* (Family Trichocomaceae). The microbial communities at SE also intermittently contained a population of yeast most closely related to the genera *Saccharomyces* and *Wickerhamomyces* (Family Saccharomycetaceae) (Figure 1b). A summary of all detected taxa can be found in Table S1.

Ordination based on Principal Coordinates Analysis (PCoA) using the weighted UniFrac distance between microbial communities including both fuel and biofilms indicated that these communities differed by site (Figure 2). Eukaryotic samples from SW were more similar to one another than samples from SE were to one another (Figure 2b). Small differences were detected between samples of fuel and biofilm at each sampling point among the bacteria (Supplemental figure S4a), but there was little to no difference among the eukarya (Supplemental figure S4b).

**Figure 2.**
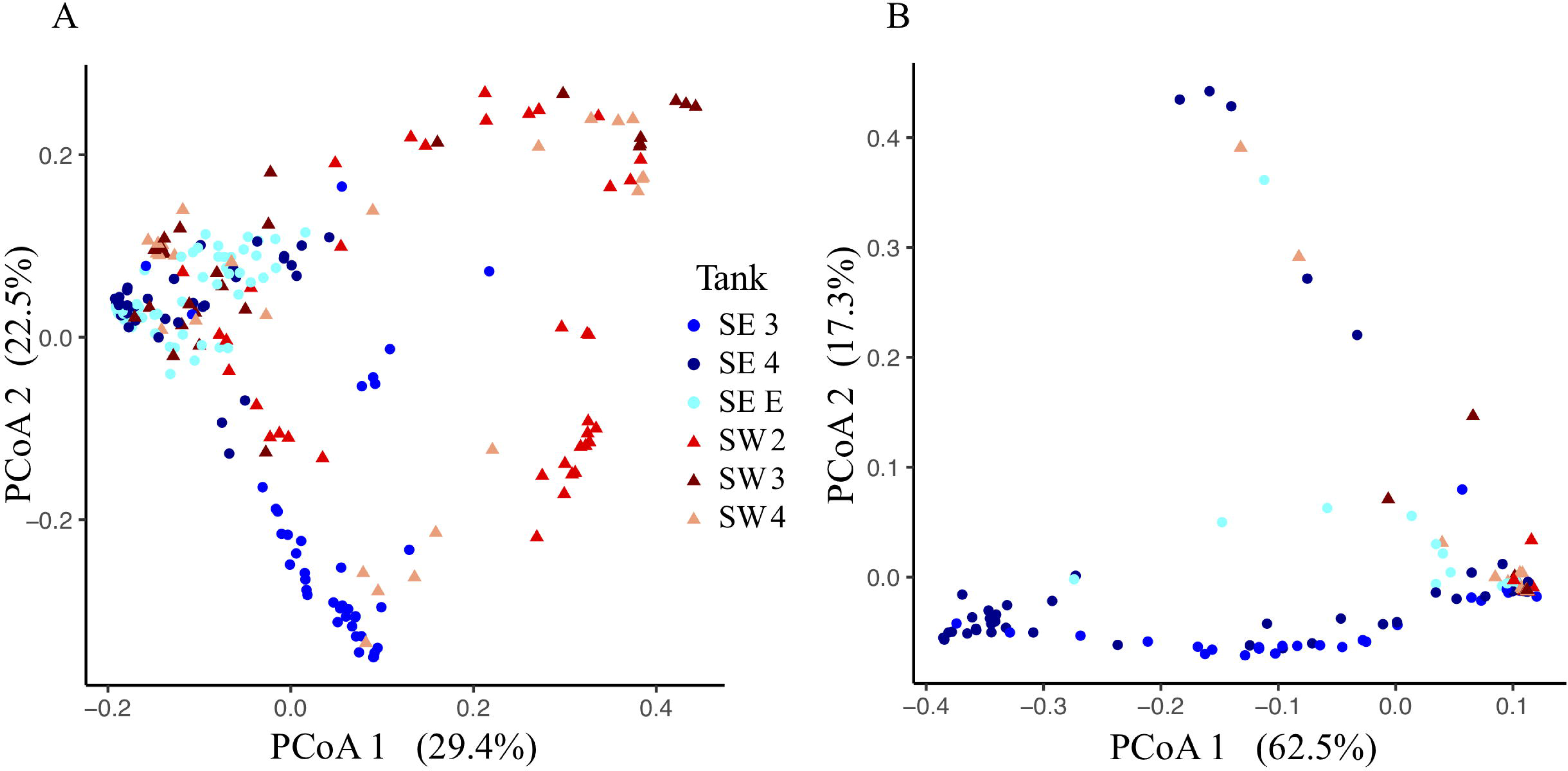
Weighted UniFrac principal component ordination of the Bacterial (A) and Eukaryotic (B) biofilm communities. Points are separated by location-circles for SE and triangles for SW. Colors represent individual tanks at each location.

### Site Observations, Biomass, and Corrosion Results

Total dry weight biomass after 12 months of exposure to fouled fuel was greatest at SW, with SW 2 having the greatest fouling of coupon surfaces (Figure S5). Overall, coupons at SW were consistently coated in greater amounts of biofilm/material (Figure S6) and had greater final biomass than those sampled at SE.

Witness coupons from tanks at SW consistently experienced greater amounts of corrosion than those at SE (Figure S7). All coupons exposed to SW tank systems had significantly more corrosion than controls exposed to sterile fuel, while SE 4 coupons were not significantly different from sterile fuel controls (Supplemental table S2). Coupons exposed to conditions in SW USTs had deeper pits than coupons from the SE location and controls (Figure 3). Over time, the skewedness of pit distribution on exposed UST coupons was greater than the controls (Figure 3). Surface roughness also increased over time (Supplemental figure S8) and was the greatest on coupons from SW 2. Tanks at SW, and SE 3 had the greatest maximal pitting rate (Table 2), with SW 2 and 4 bottom coupons having rates exceeding 8 MPY. Coupons placed at the bottom of tanks had deeper pits (Figure 3, p < 0.001), but surface roughness was not significantly different between coupons exposed to the top or bottom of the tank (p = 0.34, figure S4). There was little to no effect on O-ring load strength, tensile strength, or elongation among tanks (Supplemental figure S8).

**Table 2.**
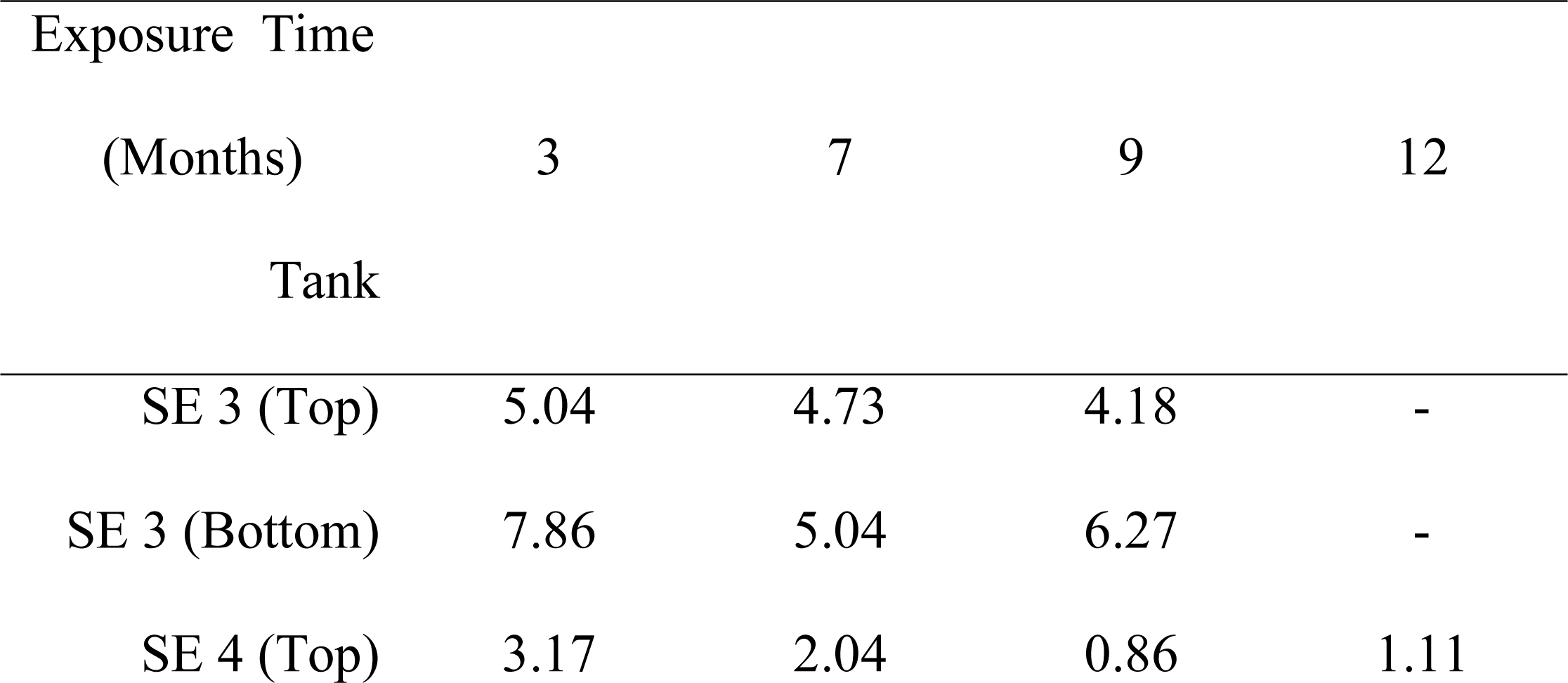
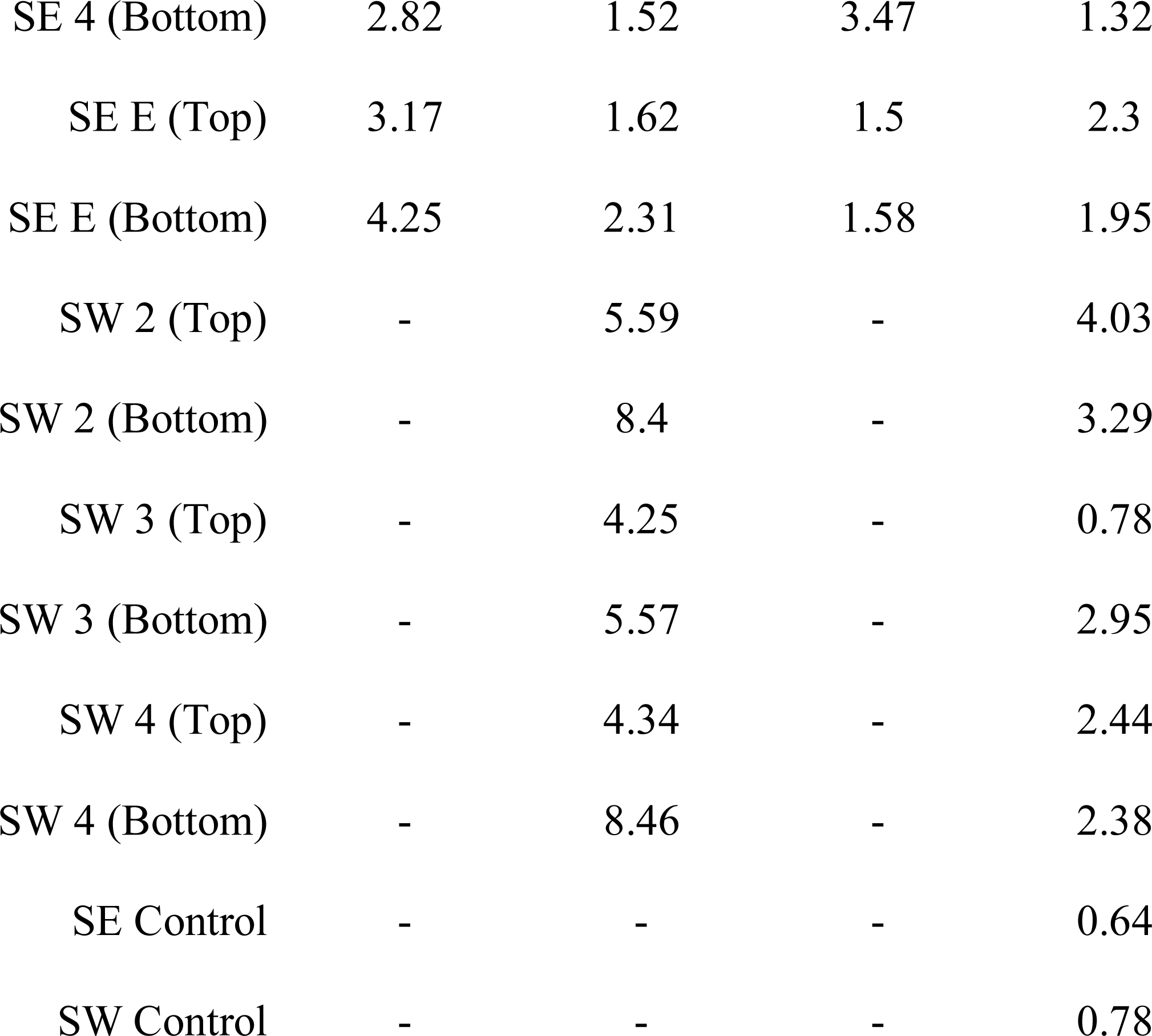
Maximum pitting rate in mils per year (MPY) calculated from the deepest pit observed in each tank, at each time point.

| Exposure Time |  |  |  |  |
| --- | --- | --- | --- | --- |
| (Months) | 3 | 7 | 9 | 12 |
| Tank |  |  |  |  |
| SE 3 (Top) | 5.04 | 4.73 | 4.18 | - |
| SE 3 (Bottom) | 7.86 | 5.04 | 6.27 | - |
| SE 4 (Top) | 3.17 | 2.04 | 0.86 | 1.11 |
| SE 4 (Bottom) | 2.82 | 1.52 | 3.47 | 1.32 |
| SE E (Top) | 3.17 | 1.62 | 1.5 | 2.3 |
| SE E (Bottom) | 4.25 | 2.31 | 1.58 | 1.95 |
| SW 2 (Top) | - | 5.59 | - | 4.03 |
| SW 2 (Bottom) | - | 8.4 | - | 3.29 |
| SW 3 (Top) | - | 4.25 | - | 0.78 |
| SW 3 (Bottom) | - | 5.57 | - | 2.95 |
| SW 4 (Top) | - | 4.34 | - | 2.44 |
| SW 4 (Bottom) | - | 8.46 | - | 2.38 |
| SE Control | - | - | - | 0.64 |
| SW Control | - | - | - | 0.78 |

**Figure 3.**
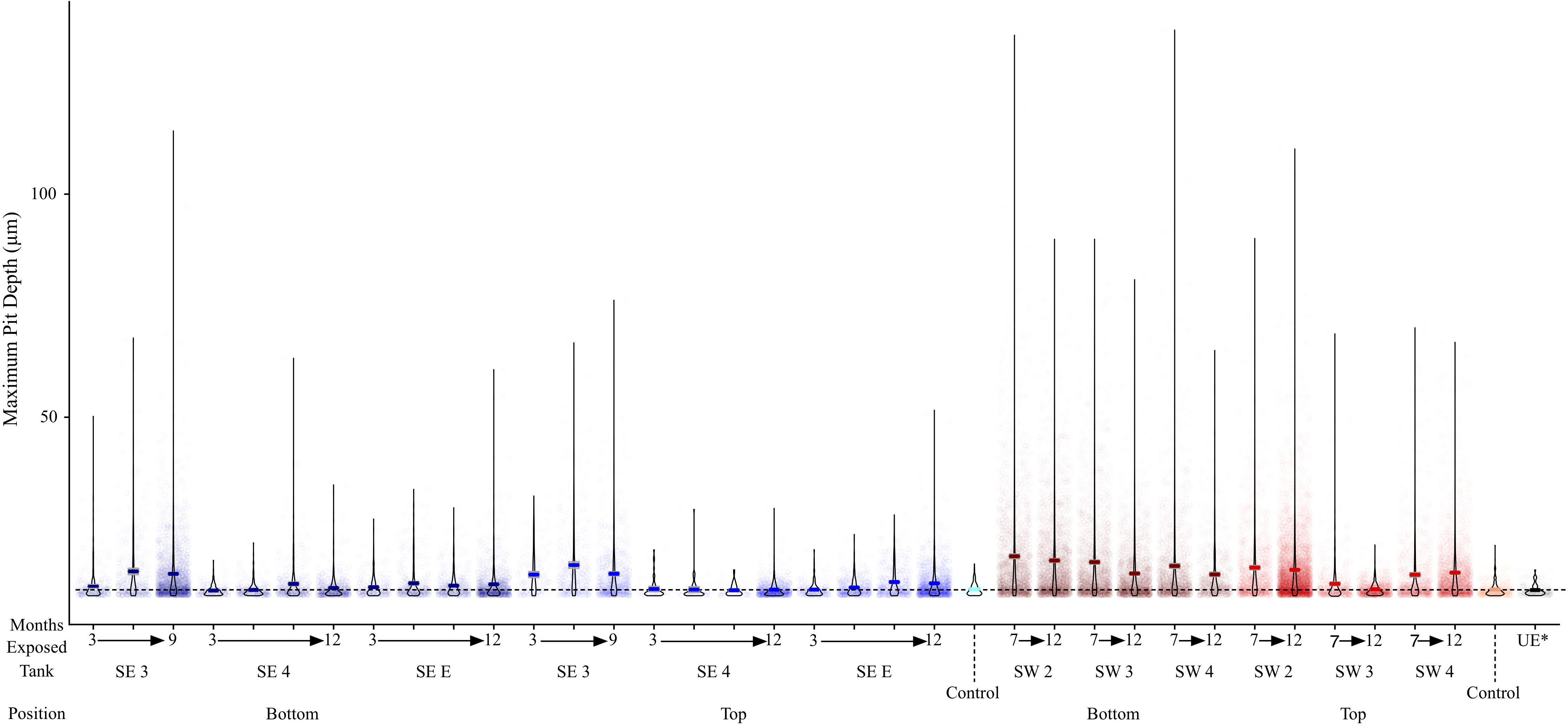
Mass loss obtained over time for each tank at SE (Blue) and SW (Red). Mean values are shown as a dark line for each sample. The dashed line represents the mean of witness coupons not exposed to fuel or field conditions.

The fuel acid number of samples taken from SE 3 increased over time from 0.17 to 1.51 mg KOH / g B20, corresponding to an elevated corrosion and maximum pit depth in SE tank 3 relative to other tanks at SE. By 9 months, fuel in SE tank 3 exceeded the ASTM standard limit of 0.3 mg KOH/g B20 for fuel acidity (Table 1). Witness coupons imaged by SEM from both bases were covered by morphologically similar biofilms, which were primarily comprised of connected filaments that are likely fungal hyphae (Figure 4c). Pits were correlated with areas that contained biofilm prior to cleaning when coupons exposed in tanks at the SE location for 3 and 7 months were imaged (Figure 4b, 4d).

**Figure 4.**
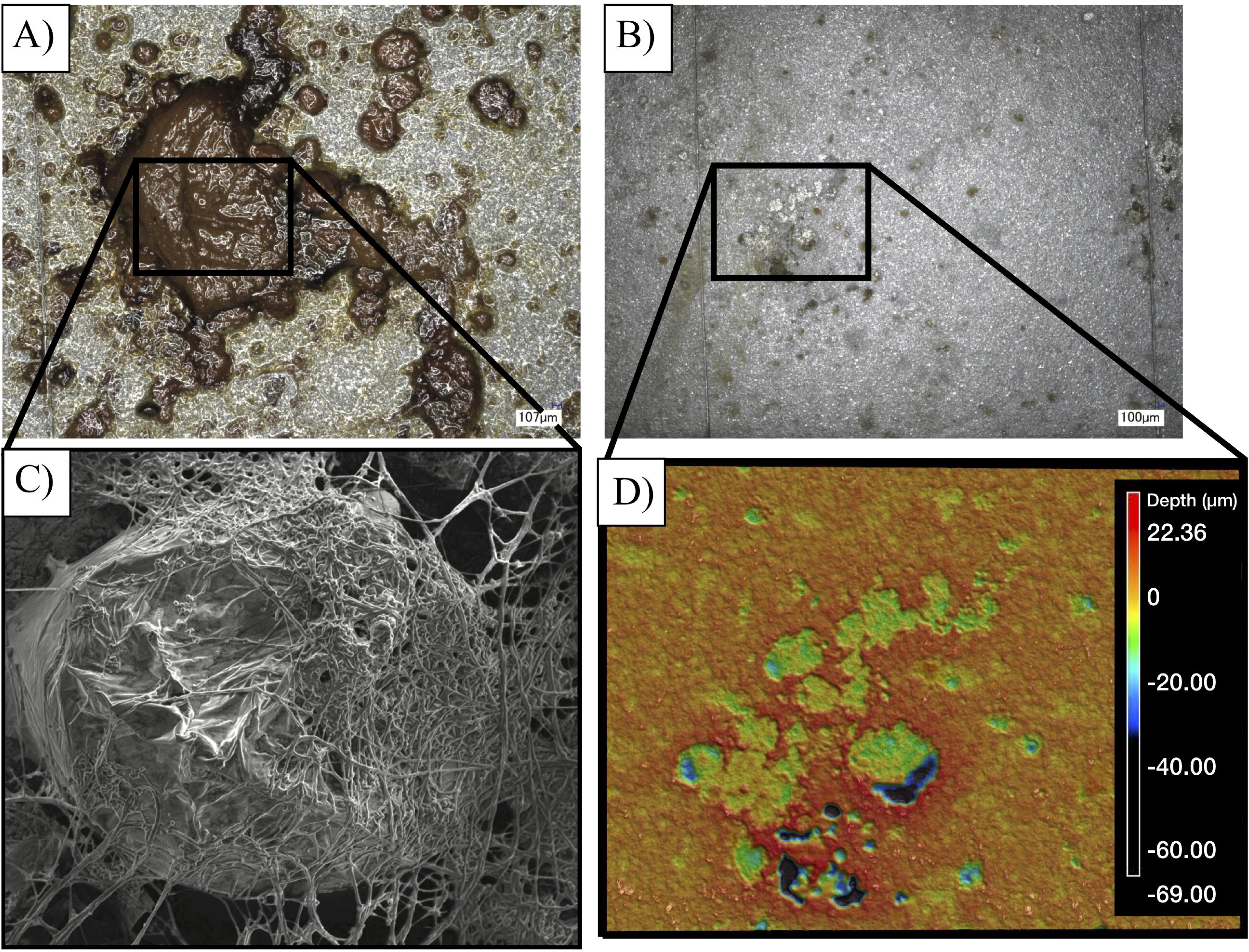
Images of a representative biofilm (A) on a witness coupon. After cleaning of the same biofilm, pitting corrosion is evident underneath the biofilm (B). SEM imaging reveals a large mass of fungal hyphae (C), while profilometry shows clear pitting corrosion underneath the fungal biofilm (D).

## Discussion

Diesel is a globally used liquid transportation fuel and is the primary ground-transportation fuel within the largest American consumer of petroleum products, the US Air Force^4^. Within the USAF, the practice of blending or replacing ULSD with a renewable fuel such as biodiesel increases fuel performance and significantly offsets carbon emissions worldwide^3^. The USAF is not alone in its increasing use of biodiesel. The use of biodiesel is on the rise, increasing worldwide from 14 thousand barrels per day (tbpd) in 2000 to 530 tbpd in 2014 (Source: U.S. Energy Administration (2015)). As usage of biodiesel and biodiesel blends increases, there is concern that increased risk of fouling or corrosion will follow. The ability of microorganisms to degrade and foul biodiesel has been reported anecdotally in operational fuel storage systems, and the link to corrosion is well studied under laboratory conditions^15,25^. Corrosion in ULSD USTs is known, with 83 percent of inspected tanks containing moderate or severe corrosion^43^ yet until now little quantifiable corrosion or microbial community data were available in operational B20 biodiesel storage tanks. Our access to in-service B20 biodiesel storage tanks was unparalleled, allowing us to carry out a temporal study of both microbial diversity and corrosion. In this work, we have provided the first *in-situ* study combining corrosion/fouling measures alongside a survey of the total microbial community in active B20 biodiesel USTs. Our study established a link between fuel fouling and increased biofilm biomass comprised of fungi and bacteria. This biomass was also linked to pitting corrosion co-located with the microbial biofilms. The linkage between fuel fouling and corrosion is of considerable concern to continuing tank operations.

There is no lack of oxidizable substrate within a tank storing B20 that supports microbial growth. Instead, mitigation of bacterial and fungal growth might be accomplished by limiting nitrogen, phosphorus, electron acceptor, or physical space near the fuel-water interface. While they were not the most abundant community members across all sampled time points, OTUs related to *Nitrospirillum* and *Burkholderia* were present within the sampled USTs. These OTUs have the capacity for nitrogen fixation and thus, could be providing the community with a source of fixed nitrogen for growth^44,45^, living in coexistence with the microorganisms most responsible for fuel acidification and corrosion. As oxygen is depleted in water near the bottom of a UST, anaerobic growth via fermentation may also provide a source of organic acids. Members of the Firmicutes genus *Caproiciproducens* can produce numerous organic acids including caproic, butyric, and acetic acid under anaerobic fermentative growth^46^ and likely contribute to increased acidity within fuel. Anaerobic microorganisms like the Clostridiaceae (including *Caproiciproducens*), were most abundant and consistently detected at SW in all three tanks. Aerobic microorganisms capable of producing acid were also present across the study, including OTUs within the Alphaproteobacteria at SW most closely related to *Gluconacetobacter* and unclassified Acetobacteraceae. Gluconacetobacter and Acetobacteraceae can produce large quantities of acetic acid in addition to producing viscous biofilms of bacterial cellulose that may enhance their ability to maintain a physical presence at the fuel-water interface as well as exacerbate both corrosion and fouling^47,48^.

In contrast to the bacteria, the fungal community, representing the bulk of detected Eukarya, was less diverse and more homogenous across both sampled locations. A single, highly abundant OTU most closely related to *Byssochlamys* was found across all tanks exhibiting fouling*. Byssochlamys* is more commonly associated with contamination of acidic fruit juices and is capable of propagating across a broad range of pH and temperature^49^. Recently, however, it was isolated from biodiesel in Hawaii, and characterized for its ability to degrade ULSD and biodiesel^50^. *Byssochlamys* is likely well adapted to grow within B20 biodiesel USTs. Members of *Byssochlamys* can withstand boiling (high temperatures are a part of the biodiesel production process), low concentrations of oxygen that may exist at the bottom of a fuel tank, and broad changes in pH as microbial community members begin to acidify water near the bottom of a fuel tank. It is also likely that these properties give *Byssochlyamys* a competitive advantage within USTs; relative abundance within the Eukarya was mostly unchanged over time with much lower diversity than the Bacteria. Filamentous fungi such as *Byssochlamys* may also provide a surface for bacteria to exploit, encouraging biofilm production within a UST^51^. SEM analysis of fouled coupons showed co-location of rod-shaped bacteria within the fungal filaments (data not shown). Ours is the first study to conclusively show the progression of biocontamination dominated by *Byssochlamys-*like OTUs in B20 biodiesel *in situ,* and to establish the relationship between their abundance and *in situ* corrosion.

The observed corrosion likely resulted from the presence of biofilms and microorganisms. The localized production of acids within biofilms can create an environment that is favorable for the induction of pitting corrosion^52^. Similar to work previously reported by Lee (2010), our *in-situ* study showed greater corrosion near the bottom of each tank where the presence of water and a fuel water interface is more likely. The interaction of fuel and water allows for enhanced microbial growth, and in turn, greater corrosion. Indeed, the greatest corrosion rates observed were in SW (> 8 MPY), where fungal fouling was the most prolific. Rates appeared to decline from 7 to 12 months, suggesting some amount of passivation may take place or an increase in uniform corrosion which could make pits appear shallower. Compared to apparent steel corrosion, polymer degradation was almost nonexistent. Fungi and Bacteria are known to degrade polymers yet no significant degradation was observed^53,54^. It is possible that the polymers tested in this study were recalcitrant to degradation over the time period of the study. Future work could include exposing polymers to contaminated fuel for an increased amount of time to confirm these findings.

Significant differences in community membership were detected between locations and even individual tanks. However, the correlation with measured environmental or operational variables was low. As a result, we still do not have definitive evidence as to why specific microorganisms, including both fungi and bacteria, were associated with a single tank or location. Tank material almost certainly was a factor-steel tanks represent a more reducing environment than fiberglass tanks, encouraging the growth of anaerobic microorganisms such as *Caproiciproducens* as observed in SW. Otherwise, stochastic forces may determine which organisms are first introduced to each tank, and the broadly shared metabolism of FAME degradation could allow subsequent growth of a broad diversity of microorganisms. As the organic acid by-products of FAME degradation accumulate, the acidity of the fuel and water would increase, selecting for acid tolerant organisms like *Byssochlamys* or *Gluconacetobacter*^47,49^, which most likely represent the climax community of fouled biodiesel. Through the process of fouling biodiesel, the ability of fungi to withstand large changes in pH and oxygen concentration almost certainly allows them to dominate the biodiesel tank ecosystem.

Tanks at SW were cleaned just prior to the beginning of *in-situ* measurements, and within 6 months, a dense fungal mat was visible across many of the USTs. Biofilms were repeatedly found on monitoring equipment and tank surfaces near the bottom of each tank. Microbial fouling of monitoring equipment prevents accurate monitoring of fuel or water levels within each tank^56^ and increases maintenance costs. The extent to which fuel acidification damages downstream vehicular fuel systems and engines is currently unknown and should be the focus of further study^57^.

The work presented here, among several other laboratory-based studies^13,58,59^ illustrates the susceptibility of biodiesel and its blends to microbial proliferation (or fouling), biodegradation, and MIC of associated infrastructure. By specification, ULSD can contain up to 5 % biodiesel by volume in the United States^60^. The worldwide use of biofuels, including biodiesel blends such as B20, will almost certainly continue. Instead of eliminating the use of biodiesel, the negative consequences of biodiesel use can be neutralized by proactive monitoring and mitigation of B20 biodiesel storage systems. Future monitoring should include sampling of fuel/water at the bottom of a UST to determine if microorganisms are present. Large scale decontamination of aircraft or buildings is possible^61^. In addition to preventative or ‘shock’ biocide treatments, sterilization technologies could be adapted to underground tank storage systems^62^ to limit biomass accumulation during normal operation. While B20 presents new storage challenges to operators, risk assessments informed by this study will aid each operator in formulating the appropriate response if contamination is detected.

## Conclusions

A mixed microbial community of filamentous fungi and acid-producing bacteria were responsible for steel corrosion in B20 biodiesel USTs. This research extends beyond *in-situ* surveys, and work is underway to characterize the most abundant fungal member of fuels and biofilms observed at both locations^63^. Biodiesel use continues to increase worldwide (Source: U.S. Energy Administration (2015)). We successfully investigated the microbial diversity at multiple USTs in two geographically distinct locations, and importantly, linked corrosion to the presence of microorganisms. The synthesis of multiple lines of evidence collected from witness samples, including microscopic characterization of biofilm communities and coupon surfaces, and measurements of corrosion, and microbial diversity, also suggests that biodiesel blends are subject to fouling by bacteria and largely, fungi. Microbial contamination and proliferation in biodiesel poses a risk to fuel storage infrastructure worldwide.

## AUTHOR INFORMATION

### Author Contributions

BWS carried out sampling, experimentation, and wrote the manuscript. CLB, CAD, JGF, HSN, carried out sampling, experimentation, and edited the manuscript. PFL, KAB, and ARN carried out experiments and edited the manuscript. WCG and BSS carried out sampling, edited the manuscript, and conceived of the experiments. All authors have given approval to the final version of the manuscript.

### Funding Sources

The work presented was supported by the United States Air Force, AFRL Biological Materials and Processing Research Team, Materials and Manufacturing Directorate as a subcontract to BSS (S-111-016-001) through UES’s prime contract FA8650-15-D-5405, task order 001; and partly through a grant awarded to BSS from the United States Air Force Academy through the Secretary of Defense’s Corrosion Protection Office Technical Corrosion Collaboration program (TCC; FA7000-15-2-0001).

#### ACKNOWLEDGMENTS

We wish to acknowledge the work and dedication of the men and women of the US Air Force who were critical in facilitating sampling at both locations. We especially thank Mr. William E. Koff, Jr. who provided invaluable institutional wisdom and experience, facilitated sampling, provided support personnel, and coordinated all logistical support. The manuscript was posted as a preprint under the accession https://doi.org/10.1101/399428^64^. Portions of the data and analysis were a part of the lead author’s dissertation^65^. Manuscript approved for public release on 21 Aug 2018, case number 88ABW-2018-4128.

## ABBREVIATIONS

ULSD: Ultra Low Sulfur Diesel
B20: 20 percent blend of biodiesel and ULSD
FAME: Fatty Acid Methyl Ester
MIC: Microbiologically Influenced Corrosion
USAF: US Air Force
DoD: Department of Defense
FY: Fiscal Year
SE: Southeast
SW: Southwest
OTU: operational taxonomic unit

## Supporting Information

**Figure S1.** Schematic representation of a B20 storage tank. Samples were taken from a “manway” access point (1), and witness coupons were suspended by chain near the bottom of the tank (2), to attempt to expose materials to fuel, as well as to any potential water bottom (3). Other potential ingress points to the tank include the fuel inlet (4) and sampling port (5).

**Figure S2.** Image of a representative coupon sampling rig placed with each tank, prior to exposure.

**Figure S3.** Overview workflow for environmental sampling of all sample types, including materials testing of o-Rings (left), uncoated carbon steel (right), and all samples destined for DNA extraction (center).

**Figure S4.** Comparison of the Bacterial (A) and Eukaryotic (B) microbial communities of both fuels and biofilm samples taken at each time point, for each tank.

**Figure S5.** Biomass obtained from coated witness coupons from both SE and SW after one year of exposure within each tank.

**Figure S6.** Images of coupons after removal from tanks at 3, 7, and 9 months (SE), or 7 months (SW).

**Figure S7.** Maximum pit depth measured for uncoated steel witness coupons from SE (Blue) and SW (Red). Mean values are shown as a dark line for each sample. The dashed line represents the mean of witness coupons not exposed to fuel or field conditions.

**Figure S8.** Roughness (S_a_) values of uncoated steel witness coupons from SE (Blue) and SW (Red).

**Figure S9.** O-Ring measurements of load (A), tensile strength (B), and elongation (C) after exposure to fuels at SE (Blue) and SW (Red). Controls exposed to SE fuel are shown in grey, and unexposed controls are represented in black.

**Table S1.** Summary statistics of Bacterial/Archaeal and Eukaryotic small subunit ribosomal RNA gene sequencing libraries.

**Table S2.** Pairwise significance tests of corrosion both between tanks and between tanks subset into individual time points.

